# Predicting Immune Escape with Pretrained Protein Language Model Embeddings

**DOI:** 10.1101/2022.11.30.518466

**Authors:** Kyle Swanson, Howard Chang, James Zou

## Abstract

Assessing the severity of new pathogenic variants requires an understanding of which mutations enable escape of the human immune response. Even single point mutations to an antigen can cause immune escape and infection by disrupting antibody binding. Recent work has modeled the effect of single point mutations on proteins by leveraging the information contained in large-scale, pretrained protein language models (PLMs). PLMs are often applied in a zero-shot setting, where the effect of each mutation is predicted based on the output of the language model with no additional training. However, this approach cannot appropriately model immune escape, which involves the interaction of two proteins—antibody and antigen— instead of one protein and requires making different predictions for the same antigenic mutation in response to different antibodies. Here, we explore several methods for predicting immune escape by building models on top of embeddings from PLMs. We evaluate our methods on a SARS-CoV-2 deep mutational scanning dataset and show that our embedding-based methods significantly outperform zero-shot methods, which have almost no predictive power. We also highlight insights gained into how best to use embeddings from PLMs to predict escape. Despite these promising results, simple statistical and machine learning baseline models that do not use pretraining perform comparably, showing that computationally expensive pretraining approaches may not be beneficial for escape prediction. Furthermore, all models perform relatively poorly, indicating that future work is necessary to improve escape prediction with or without pretrained embeddings^1^.

## 1 Introduction

Pathogens are constantly evolving in their search to evade the immune system and infect host organisms [1]. In many organisms, including humans, this evolutionary battle occurs in the context of antibody-antigen interactions [2]. Antibodies are proteins produced by the immune system that are designed to bind to antigens, which are pathogenic proteins that induce an immune response. Antibodies that effectively bind to an antigen and neutralize the pathogen put evolutionary pressure on the pathogen to mutate its antigen in a process known as immune escape [3]. Predicting which mutations cause escape is crucial to identifying dangerous pathogenic variants that can cause infection and disease even in the presence of antibodies from prior infection, vaccination, or therapies [3–5]. Machine learning models have been developed that can predict the effect of protein mutations on various protein functions [4, 6–9]. Recent approaches to mutation effect prediction have leveraged large protein language models (PLMs) that have been trained in an unsupervised manner on huge databases with hundreds of millions to billions of protein sequences [10, 11]. PLMs learn the underlying statistics of naturally occurring protein sequences and can predict the likelihood that a given amino acid appears at a position in a protein. Prior work has shown that the relative likelihoods of a mutated and wildtype amino acid at a given position are predictive of the effect of that mutation in a zero-shot manner (i.e., without additional training) [6, 7, 12, 13].

However, a major limitation of the zero-shot likelihood approach is that it predicts the same likelihood for a mutation regardless of the protein function in question [7]. Since proteins can have multiple functions that are affected differently by the same mutation, one likelihood cannot model the effect of a mutation on all of these functions simultaneously. Additionally, the likelihood only accounts for the protein that is mutated, which means that it ignores any interacting proteins such as antibodies.

We propose to overcome these limitations by modeling immune escape using antibody and antigen embeddings produced by a PLM. These embeddings encode information about the protein, including aspects of 3D structure, that can inform the effect of protein mutations [14]. We build a lightweight neural model that learns to extract information from the embeddings to predict escape in an antibody-dependent manner. We develop several variants of this embedding-based approach and evaluate them on a SARS-CoV-2 deep mutational scanning dataset from Cao et al. [5]. We show that embeddings significantly outperform zero-shot likelihoods, which have almost no predictive power. We discuss insights gained from our experiments about how best to use embeddings from PLMs to predict escape. We also develop two statistical baseline models and a machine learning model that do not rely on the pretrained models. These models perform comparably to the embedding models, indicating that pretrained embeddings may not be beneficial for predicting escape. Furthermore, the relatively poor performance of all models demonstrates that future work is necessary to improve escape prediction with or without pretrained embeddings.

## 2 Methods

Our goal is to design a model that can predict immune escape. Specifically, we model escape by predicting how single point mutations to an antigen affect the binding ability of antibodies. Below, we first outline notation used throughout this section. Then, we describe several models to predict immune escape using either simple statistics (mutation and site models), a machine learning model trained from scratch (RNN), PLM likelihoods, or PLM embeddings.

### 2.1 Notation

An antigen is a sequence of amino acids *A* = {*A*_1_, *A*_2_, …, *A*_*n*_} with each *A*_*i*_ ∈ **P** where **P** is the set of 20 naturally occurring amino acids. Each location *s* ∈ {1, 2, …, *n*} in the antigen is called a site. The original, unmutated sequence of antigen amino acids is referred to as the wildtype sequence. Due to the abundance of single point mutation data and the relative lack of multi-mutation data, we only consider single point mutations, where a single site *s* in the antigen has its amino acid mutated from the wildtype amino acid *A*_*s*_ to the mutant amino acid *M* ∈ **P \** *A*_*s*_, which is one of the other 19 possible amino acids. The mutated antigen sequence then becomes *A*^*s*→*M*^ = {*A*_1_, …, *A*_*s*−1_, *M, A*_*s*+1_, …, *A*_*n*_}. In this paper, we consider a single antigen *A* and all possible *n* × 19 single point mutations across the *n* antigen sites.

An antibody *B* is a protein that consists of four chains, where each chain is a sequence of amino acids. Among the four chains, two are identical chains called the heavy chain with the sequence *H* = {*H*_1_, *H*_2_, …, *H*_*h*_} and two are identical chains called the light chain with the sequence *L* = {*L*_1_, *L*_2_, …, *L*_*l*_}. The antibody as a whole is represented by the pair of unique chains, *B* = (*H, L*). Here, we consider many different antibodies *B* ∈ **B** where **B** is a set of antibodies, all of which bind to the same antigen *A*.

When an antibody comes into contact with an antigen, interactions between the amino acids of the antibody and antigen can result in binding and subsequent neutralization of the pathogen. Mutations to the antigen may inhibit antibody binding. The degree to which antibody binding is reduced by an antigenic mutation is called the escape score. For antigen *A* with amino acid *M* at site *s* in the presence of antibody *B* that binds the wildtype antigen, the experimentally determined escape score is *E*(*A, s, M, B*) ≥ 0, with larger numbers indicating more escape (less antibody binding with the mutation). If *M* is the wildtype amino acid at site *s*, i.e., *A*_*s*_ = *M*, then the escape score is zero since the antigen is unchanged. If there is a mutation so that *A*_*s*_ *≠ M*, then the escape score may be zero or non-zero depending on whether and to what degree the mutation affects antibody binding.

Given a set of training data points **T** = {(*A, s, M, B*)} _*s*∈{1,2,…,*n*}, *M*∈**P**,*B*∈**B**_ consisting of one fixed antigen with many site, mutation, and antibody combinations, along with their known escape scores given by *E*, our goal is to build a model that can predict the escape score for a site, mutation, and antibody combination not in the training set.

### 2.2 Mutation Model

The mutation model `MM` models escape as a function of the change in amino acid from wildtype to mutant. This assumes that amino acid changes have consistent escape effects regardless of the site and antibody. The model is fitted by computing the average escape score in the training set for each pair of wildtype (wt) and mutant (mut) amino acids across all sites and antibodies, i.e.,

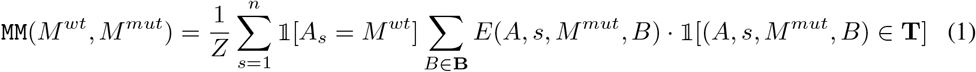

where *n* is the number of antigen sites, is an indicator variable, and

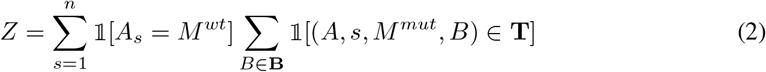

is the total number of data points in the training set where site *s* is mutated from *M*^*wt*^ to *M*^*mut*^. To make a prediction for a new site *s*, mutation *M*, and antibody *B*, the model simply outputs `MM`(*A, s, M, B*) = `MM`(*A*_*s*_, *M*), thereby ignoring the site, the rest of the antigen sequence, and the antibody sequences. Since there are 20 amino acids, this model has 20 × 20 = 400 parameters.

### 2.3 Site Model

The site model `SM` models escape as a function of the antigen site. This assumes that sites have consistent escape effects regardless of the wildtype and mutant amino acids and the antibody. The model is fitted by computing the average escape score in the training set for each antigen site across all mutant amino acids and across all antibodies, i.e.,

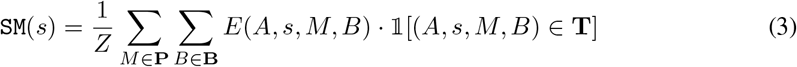

where 𝟙 is an indicator variable and

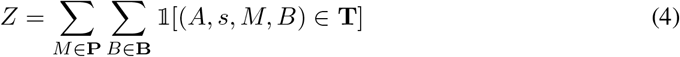

is the total number of data points in the training set for site *s*. To make a prediction for a new site *s*, mutation *M*, and antibody *B*, the model simply outputs `SM`(*A, s, M, B*) = `SM`(*s*), thereby ignoring the entire antigen sequence, including wildtype and mutant amino acids, and the antibody sequences. The model has one parameter for each antigen site for a total of *n* parameters.

### 2.4 RNN

We train a recurrent neural network (RNN) from scratch as a non-pretrained baseline machine learning model. Given antigen *A*, site *s*, and mutated amino acid *M* at that site, we construct the mutated antigen sequence *A*^*s*→*M*^ and embed it using a bidirectional LSTM [15]. We then extract an embedding in one of two ways and pass that embedding through a small multilayer perceptron to predict escape. In the model we call RNN Seq, this embedding is the final LSTM hidden state embedding representing the whole sequence. In the model we call RNN Res, this embedding is the LSTM output embedding corresponding to the mutated site *s*. Since the RNN model ignores the antibody sequences, it computes `RNN`(*A, s, M, B*) = `RNN`(*A, s, M*).

### 2.5 Likelihood Model

Our likelihood model `L` adopts the zero-shot mutation prediction framework of Meier et al. [7]. In this framework, a PLM is applied to the antigen sequence *A*^*s*→^^<mask>^, where the amino acid at site *s* is replaced with a `<mask>` token. The escape score is predicted as the model’s log odds ratio of the mutated amino acid *M* versus the wildtype amino acid *A*_*s*_ at site *s*. Specifically, the model computes

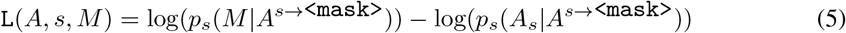

where *p*_*s*_(·|·) is the probability the model assigns to an amino acid at a site *s* within the given sequence. The likelihood model does not require any additional training. It does not incorporate the antibody sequences so `L`(*A, s, M, B*) = `L`(*A, s, M*).

### 2.6 Embedding Models

PLM embeddings provide a more flexible way of modeling immune escape than PLM likelihoods since it is possible to combine embeddings of multiple protein sequences in a single model. In our PLM embedding models, we train a small multilayer perceptron to use some form of PLM embedding as input to predict the escape score. All of the models use an embedding of the mutated antigen sequence *A*^*s*→*M*^, and some additionally use an embedding of the wildtype antigen sequence *A* and/or embeddings of the antibody heavy and light chains. The embedding variants are described below and illustrated in Figure 1.

**Figure 1:**
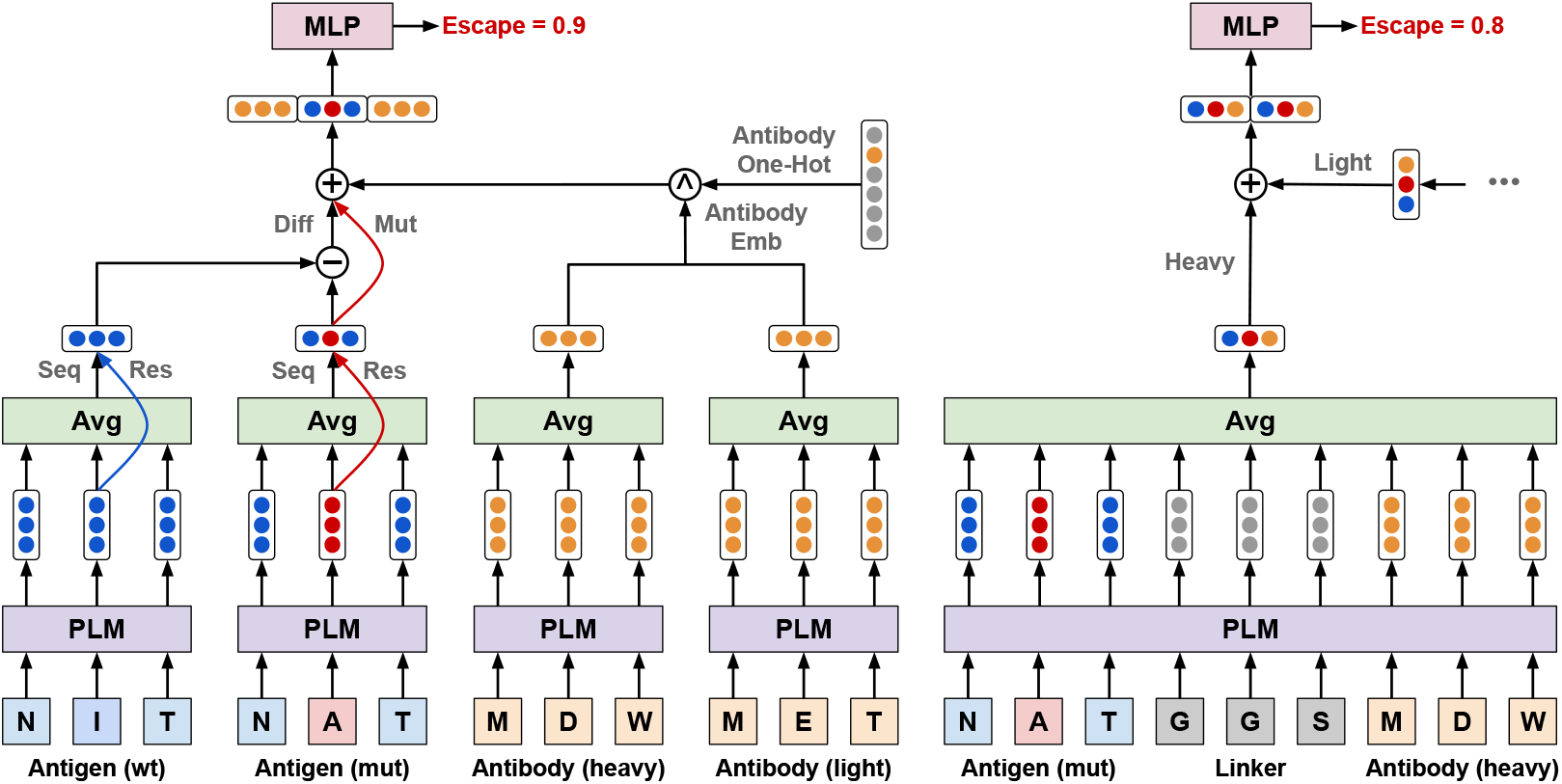
An illustration of the various PLM embedding models for predicting immune escape. The different embedding types are described in detail in Section 2.6. Note that ⊕ indicates concatenation, ⊖ indicates elementwise difference, and 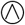 indicates exclusive or (i.e., only one vector is used).

#### Antigen Sequence Mutant

The PLM is given the mutated antigen sequence *A*^*s*→*M*^ and computes the embedding matrix *R*^*mut*^ = `PLM`(*A*^*s*→*M*^) with *R*^*mut*^ ∈ ℝ^*n*×*d*^ containing a *d*-dimensional embedding for each amino acid in the antigen sequence. The PLM embedding at each site encodes the identity of the amino acid at that site as well as its role in the context of the antigen sequence. The PLM embeddings of the amino acids are averaged to form a single embedding for the full antigen sequence, 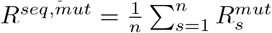 with *R*^*seq,mut*^ ∈ ℝ^*d*^. We refer to this embedding as Antigen Seq Mut.

#### Antigen Residue Mutant

As above, the PLM computes the embedding matrix *R*^*mut*^ = `PLM`(*A*^*s*→*M*^). Here, the embedding 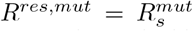 of the mutated residue at site *s* is used instead of the sequence average. We refer to this embedding as Antigen Res Mut.

#### Antigen Difference

Embeddings are computed for both the mutated antigen sequence *A*^*s*→*M*^ and the wildtype antigen sequence *A* as *R*^*mut*^ = `PLM`(*A*^*s*→*M*^) and *R*^*wt*^ = `PLM`(*A*). The difference between these embeddings is then computed, either at the sequence level or at the residue level, as *R*^*seq,diff*^ = *R*^*seq,mut*^− *R*^*seq,wt*^ or *R*^*res,diff*^ = *R*^*res,mut*^ − *R*^*res,wt*^. We refer to these embeddings as Antigen Seq Diff and Antigen Res Diff embeddings. We also concatenate the mutant and difference embeddings at either the sequence or residue level to form what we call Antigen Seq MutDiff and Antigen Res MutDiff embeddings, which are 2*d*-dimensional.

#### Antibody

We develop four methods of incorporating antibody information into the model.

1. **Antibody One-Hot:** We concatenate the antigen embedding (of any form) with a one-hot encoding of the antibody to provide the model with the antibody identity but without embedding information.
2. **Antibody Emb:** We use the PLM to embed the heavy and light chains of the antibody, and we concatenate those two embeddings with the antigen embedding to create a 3*d*-dimensional embedding.
3. **Antibody Att:** We use attention to merge antigen and antibody embeddings. Due to slow training and poor performance, we reserve this model’s description and results for Appendices C and D.
4. **Antigen Linker Antibody:** We create embeddings of combined antibody-antigen sequences. For each antibody, we create two sequences, both including the mutated antigen *A*^*s*→*M*^ and one with the heavy chain *H* and the other with the light chain *L*. In both cases, the antigen and antibody sequences are joined by seven repeats of a glycine-glycine-serine linker. Both linked sequences are embedded by the PLM, sequence averaged, and then concatenated to form a single 2*d*-dimensional embedding for the antigen and antibody. This design is inspired by the use of linkers to enable protein complex prediction from single-chain protein structure prediction models like AlphaFold2 [16].

## 3 Experiments

Here, we describe the data we use to train and evaluate our model as well as the data splits, tasks, metrics, and models that we use.

### 3.1 Data

We use SARS-CoV-2 deep mutational scanning data from Cao et al. [5]. This data consists of 247 antibodies that are known to bind the original strain of SARS-CoV-2 by binding to the receptor binding domain (RBD) of the spike protein. The binding ability of each antibody is measured for the wildtype RBD antigen as well as for all 3,819 single point mutations to the antigen (201 sites in the RBD with 19 amino acid substitutions at each site). For each antibody and each antigen mutation, an escape score is computed as a normalized measure of the reduction in antibody binding compared to the wildtype antigen (see Figure 2). Of the 943,293 escape scores in the dataset, 30,658 (3.2%) are non-zero, all in the range (0, 1] except for 74 outliers above 1 with a max of 3.6 (see Figure S.1). Cao et al. [5] clustered the 247 antibodies into six groups based on their escape scores (see Figure S.1).

**Figure 2:**
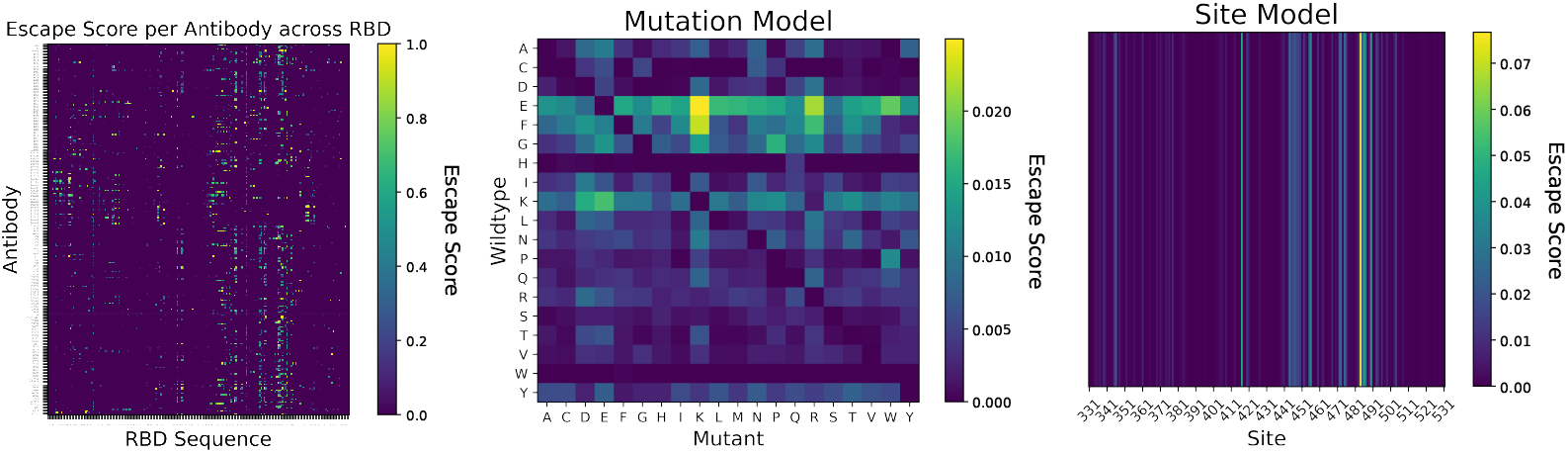
SARS-CoV-2 immune escape data from Cao et al. [5] and associated statistical models. (Left) The average escape score across all amino acid mutations for each antibody and each antigen site in the receptor binding domain (RBD) of the SARS-CoV-2 spike protein. (Middle) A mutation model fit on the full dataset, showing the escape score for each wildtype to mutant amino acid change averaged across all antigen sites and antibodies. (Right) A site model fit on the full dataset, showing the escape score for each antigen site averaged across all amino acid mutations and antibodies. Note: In all figures, the 74 escape scores greater than 1 (max 3.6) are truncated to 1.

### 3.2 Data Splits

The practical usefulness of an escape prediction model, as well as the difficulty of learning such a model, depends on how the data is split. Below we describe and motivate the data splits we use.

#### Mutation

Mutations are randomly split between train and test. This assumes that for a new antibody, we already know escape scores for some but not all mutations across all antigen sites. This split corresponds to a scenario in which we have a significant amount of escape data, either from laboratory mutation experiments or real-world infections by mutated pathogens, across antigen sites for a particular antibody. However, we do not have escape data for the complete set of mutations, so a model trained in this setting would be able to fill in the escape effect of any missing mutations.

#### Site

Antigen sites are randomly split between train and test. This assumes that for a new antibody, we already know escape scores for some but not all antigen sites. In this split, we are more conservative and assume that we only have escape data for some antigen sites and need to make predictions for other antigen sites. This still requires knowing some escape scores for the antibody, but we no longer need to know escape scores across all sites, making it possible to use any available escape data.

#### Antibody

Antibodies are randomly split between train and test. This assumes that we do not know any escape scores for a new antibody. This split models a situation in which some antibodies have already been experimentally evaluated and have escape data, and we want to make predictions for a new antibody for which we do not yet have any escape data.

#### Antibody group

Antibody groups, as defined by a clustering of escape scores, are randomly split between train and test. This assumes that we do not know any escape scores for a new antibody, and furthermore, no antibody in the train set has a similar pattern of escape to this antibody. This split is especially relevant since new groups of antibodies may continue to bind the antigen and eliminate the pathogen even in the presence of antigenic mutations that prevent binding to other antibody groups.

In general, the antibody and antibody group splits are more practically useful because they demonstrate the effectiveness of escape prediction for antibodies that have not undergone any experimental escape measurements. This means that new antibodies can be evaluated entirely *in silico*. Escape prediction models that are effective under these data splits could thus be used to guide the selection or design of antibodies that are robust to antigenic mutations that escape other antibodies, providing an avenue for designing effective new antibody treatments against mutating pathogens.

For all four splits, we train and test the models across all antibodies (cross-antibody setting) using five-fold cross-validation. For the mutation and site splits where each antibody can appear in both the train and test sets, we also build separate models for each of the 247 antibodies (per-antibody setting). This makes it possible to compare the ability of a single model learned across antibodies to separate models learned for each antibody individually.

### 3.3 Tasks and Metrics

For all of the models except for the likelihood model, which doesn’t require training, we train the model either for a regression task, where escape scores are real values, or for a classification task, where escape scores are binarized into zero or non-zero escape. All models are evaluated with the metrics ROC-AUC (area under the receiver operating characteristic curve) and PRC-AUC (area under the precision-recall curve), and regression models are additionally evaluated with the metrics MSE (mean squared error) and R^2^ (coefficient of determination).

### 3.4 Models

Below we describe the implementation details of the models we developed. All models were built using PyTorch version 1.12.1 [17].

#### 3.4.1 RNN

The RNN model is a bidirectional LSTM [15] with a hidden dimensionality of 100. The input to the RNN is the mutated antigen sequence with amino acids encoded using trainable embeddings with a dimensionality of 100. The output of the RNN is an embedding for each amino acid with a dimensionality of 200 (100 for each direction of the RNN). Either the final hidden state embedding (RNN Seq) or the output embedding of the mutated amino acid (RNN Res) is used as input to a small multilayer perceptron (see below), which makes escape predictions. The amino acid embeddings, RNN, and multilayer perceptron are trained end-to-end.

#### 3.4.2 Protein Language Model

For the likelihood and embedding models, we use the pretrained protein language model ESM2 [14]. We specifically use the `esm2_t33_650M_UR50D` version of the model consisting of 33 layers and 650M parameters that was trained on the UniRef50 database [18]. The embeddings produced by this model have a dimensionality of 1,280 and are used as fixed input to a small multilayer perceptron.

#### 3.4.3 Multilayer Perceptron

The multilayer perceptron (MLP) model that we use with the RNN and with all of the pretrained embeddings has two hidden layers with 100 neurons in each layer and ReLU activation followed by a single linear output. For classification tasks, we apply a sigmoid activation to the output.

#### 3.4.4 Training

The RNN and the embedding models were trained with mean squared error loss for regression and binary cross entropy loss for classification using the Adam optimizer [19]. Per-antibody models were trained for 50 epochs while cross-antibody models were trained for one epoch. The RNN model was trained on a single GPU, with training taking about 3 minutes for a cross-antibody model (one fold) and about 30 seconds for each per-antibody model. The embedding models were trained on a single CPU, with training taking about 1 minute for a cross-antibody model (one fold) and about 15 seconds for each per-antibody model.

## 4 Results

In this section, we highlight some of the key results from our experiments (see Figure 3). We only show classification model results since the regression models performed poorly. Additionally, since the relative ranking of models was similar between ROC-AUC and PRC-AUC but the differences in PRC-AUC scores were more noticeable, we only present PRC-AUC results. We show results for all data splits and for a subset of the models, leaving out embedding models whose performance was not insightful for space. The complete set of results across all 168 experiments is in Appendix D.

**Figure 3:**
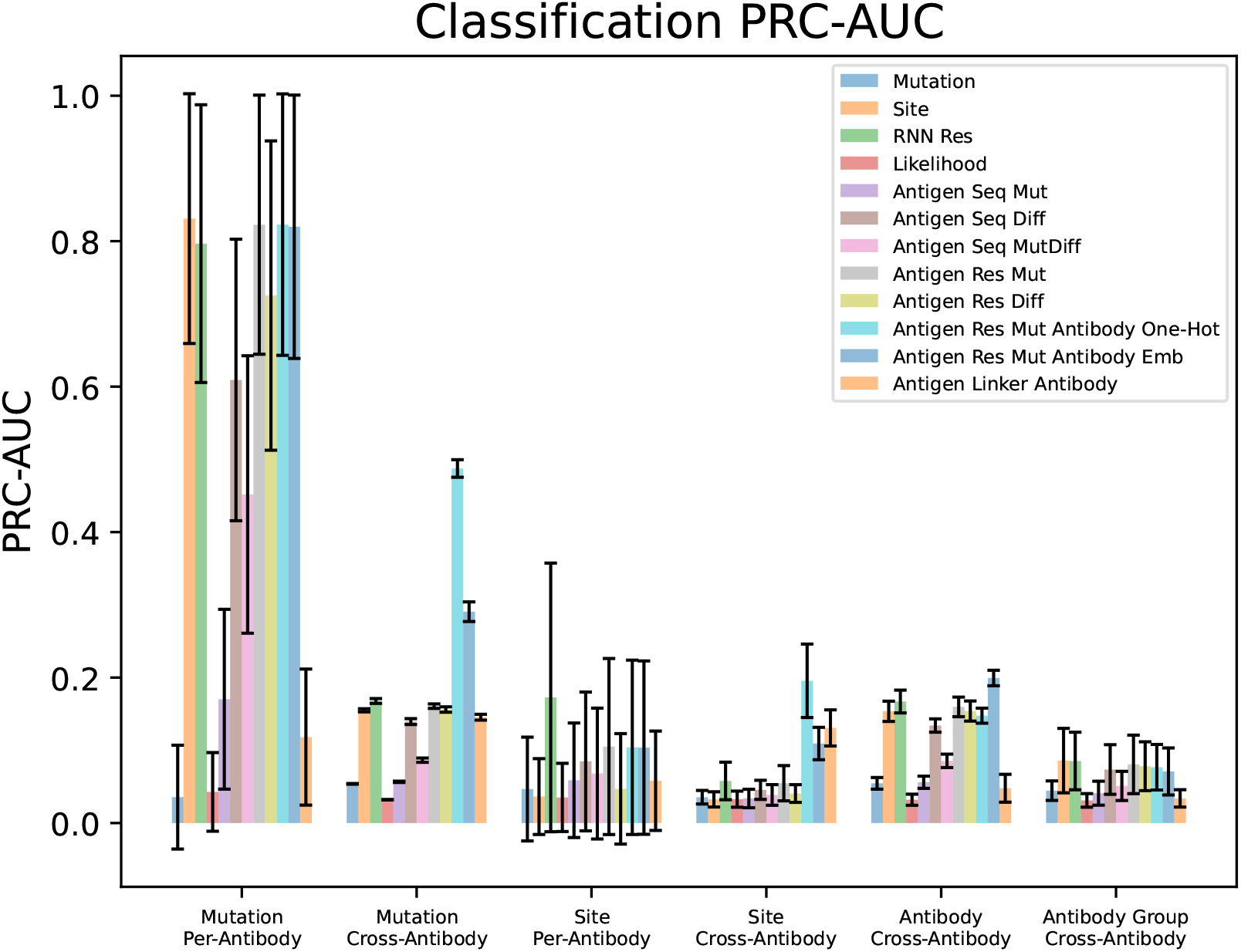
Classification model results with the PRC-AUC metric across data splits (x-axis) and models (color-coded bars). Error bars indicate the standard deviation across 247 antibodies for per-antibody splits and across five-fold cross-validation for cross-antibody splits.

### 4.1 Mutation Model

The mutation model is a very weak model. On the mutation and site splits in the per-antibody setting, the model has essentially no predictive power, and on all four splits in the cross-antibody setting, the model performs poorly. This is to be expected since the model ignores the mutation site even though the mutation site is very informative of immune escape due to the consistent interaction of key antigen sites with binding antibodies.

### 4.2 Site Model

The site model is strong across most splits with the exception of the site split where the model has no information about unseen sites. The site model is frequently competitive with the best embedding models despite containing only 201 parameters instead of 650M parameters. The site model is significantly more effective in the per-antibody mutation split than in any of the cross-antibody splits since escape is highly consistent at a given antigen site for an antibody across amino acid mutations. Even so, the fact that the model retains some predictive power across antibodies and antibody groups indicates that patterns of escape at specific sites are conserved.

### 4.3 RNN

The RNN Res model performs comparatively well across all splits. It is only outperformed by the embedding models that include antibody embeddings, which is reasonable given that the RNN only processes the antigen sequence and has no knowledge of the antibody. Notably, the model performs on par with or better than most of the embedding models, even on the site split, despite training in just a couple of minutes with no expensive pretraining needed. This shows that existing pretraining methods and models may not be particularly beneficial for escape prediction, at least for this dataset. Interestingly, the RNN Seq model performs very poorly (see Appendix D), which may indicate that sequence averaging obscures the relevant information from the mutated amino acid. A similar phenomenon occurs in the Antigen Seq versus Antigen Res embeddings, meaning that it may be preferable to use residue rather than sequence averaged embeddings across model types.

### 4.4 Likelihood Model

The likelihood model has virtually no predictive power across all data splits. This is in contrast to examples in the literature where likelihoods achieve reasonable mutation effect prediction performance [7]. This finding demonstrates a fundamental limitation of the zero-shot prediction framework since likelihoods derived from models trained to recreate naturally occurring proteins may not be calibrated to predict the probability of antigen escape.

### 4.5 Embedding Models

The embedding models significantly outperform the likelihood model across all data splits. This indicates that PLMs do contain information that is useful for mutation effect prediction but require that their representations are adapted to the task rather than used in a zero-shot manner. However, the strength of the RNN model indicates that pretrained embeddings are not necessary for escape prediction. Even so, the embedding model results still provide several interesting takeaways regarding how best to use pretrained embeddings to predict escape in cases where they may be useful.

#### Mutant vs Difference

The Antigen Seq Diff embedding consistently outperforms the Antigen Seq Mut embedding, which indicates that the change in embedding from wildtype to mutant is more informative than the mutant embedding in isolation. The concatenation of the mutant and difference embeddings (MutDiff) does not improve performance further, indicating that the mutant embeddings do not contribute information beyond that contained in the difference embeddings.

#### Sequence vs Residue

The Antigen Res Mut embedding outperforms the Antigen Seq embeddings, perhaps because the sequence embeddings contain largely irrelevant information from the non-mutated residues. Interestingly, using embedding differences (Antigen Res Diff) instead of mutant embeddings (Antigen Res Mut) does not improve performance at the residue level (see Appendix D).

#### Antibody

Including antibody information alongside the antigen embeddings generally provides a benefit in all cross-antibody splits, where each model sees more than one antibody. The Antigen Res Mut Antibody One-Hot encodings provide a particularly large benefit in the mutation and site cross-antibody splits, where the same antibody can appear in train and test and the one-hot encoding makes it easy for the model to associate patterns of escape with particular antibodies. Interestingly, in the mutation splits, the one-hot antibody encoding does not allow the cross-antibody model to recover the performance of the per-antibody models, which may suggest a benefit to training separate models for each antibody, even though each model will be trained on less data.

Although the antibody embedding in the Antigen Res Mut Antibody Emb model should also indicate the antibody identity, the model is not able to use this information as well as the one-hot embedding in the mutation and site cross-antibody splits. However, in the antibody and antibody group splits, no antibodies are shared between train and test and the one-hot encoding provides no useful information at test time. In contrast, the antibody embedding provides a small performance boost in the antibody split, but this effect disappears in the harder antibody group split where no similar antibodies are present in the test set. This indicates that the model struggles to learn how to extract useful information from the embeddings of antibodies with diverse antigen binding behavior.

The Antigen Linker Antibody embeddings do provide some benefit over antibody-agnostic models in the site cross-antibody split, but otherwise they do not help and sometimes hurt performance, as in the antibody split. This is likely because the PLM was not designed to use linkers and may not provide particularly useful embeddings for such artificially linked sequences.

## 5 Conclusion

We present several methods for predicting immune escape using pretrained protein language model embeddings. We performed a comprehensive set of experiments on a SARS-CoV-2 deep mutational scanning dataset and showed that embeddings from PLMs are much more effective at predicting escape than zero-shot likelihoods. The Antigen Res Mut Antibody Emb model was particularly powerful among the embedding models, indicating that escape is best modeled at the residue level with both antigen and antibody embeddings. Although these results are promising, the relatively strong performance of the site model and the RNN, neither of which rely on computationally expensive pretraining, show that PLM embeddings may not be particularly beneficial for tasks such as escape prediction. Furthermore, the overall poor performance of all models across most splits demonstrates that significant future work is needed to make accurate and useful escape predictions. Notably, the results here are limited to a single antibody-antigen escape prediction task and dataset. Although the comprehensive nature of this data, which includes every possible single point mutation of the antigen for every antibody, gives our conclusions strength, experimentation on additional datasets is necessary to validate whether the conclusions drawn here generalize to other escape prediction tasks.

## Acknowledgments

We would like to thank Mert Yuksekgonul, Mirac Suzgun, Jeremy Wohlwend, and Kirk Swanson for their insightful comments, suggestions, and feedback. We would also like to thank the members of the Chang lab and the Zou lab for their helpful discussions. K.S. gratefully acknowledges the support of the Knight-Hennessy Scholarship.

## A Data and Code Availability

The SARS-CoV-2 deep mutational scanning data from Cao et al. [5] is available at https://github.com/jbloomlab/SARS2_RBD_Ab_escape_maps/tree/main/data/2022_Cao_Omicron. The file `data.csv` contains the escape data for each antibody-antigen mutation combination, and the file `antibodies.csv` contains the sequences for the heavy and light chains for all the antibodies. The ESM2 pretrained protein language model [7] that we used is the `esm2_t33_650M_UR50D` model from https://github.com/facebookresearch/esm. Our code, data, embeddings, and results are available at https://github.com/swansonk14/escape_embeddings.

## B Data Visualization

In addition to Figure 2, Figures S.1 and S.2 further visualize the SARS-CoV-2 deep mutational scanning data from Cao et al. [5]. Figure S.1 shows a histogram of the non-zero escape scores. Figure S.1 shows the escape scores across antibodies and antigen sites with antibodies divided into groups according to the escape-based clustering performed by Cao et al. [5].

## C Attention Model

The antibody attention model (Antibody Att) works as follows. First, PLM embeddings are created for the mutated antigen sequence and both chains of the antibody: *R*^*mut*^ = `PLM`(*A*^*s*→*M*^), *R*^*H*^ = `PLM`(*H*), and *R*^*L*^ = `PLM`(*L*). Then, we apply multi-head attention (using PyTorch’s `MultiheadAttention` module with eight attention heads) twice, once with the antigen and antibody heavy chain embeddings and once with the antigen and antibody light chain embeddings. In both cases, the antigen embedding is used as the query while the antibody chain embedding is used as the key and value. This gives us *R*^*att,H*^ = `MultiheadAttention`(*R*^*mut*^, *R*^*H*^, *R*^*H*^) and *R*^*att,L*^ = `MultiheadAttention`(*R*^*mut*^, *R*^*L*^, *R*^*L*^) with *R*^*att,H*^ ∈ ℝ^*n*×*d*^ and *R*^*att,L*^ ∈ ℝ^*n*×*d*^ where *n* is the length of the antigen sequence and *d* is the embedding dimensionality. Next, we take the average across the antigen sequence to get 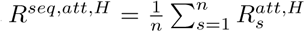 and 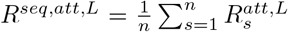 with *R*^*seq,att,H*^ ∈ ℝ^*d*^ and *R*^*seq,att,L*^ ∈ ℝ^*d*^. Finally, we concatenate *R*^*seq,att,H*^ and *R*^*seq,att,L*^ to form a 2*d*-dimensional embedding that is used as input to an MLP to predict escape. Since the attention model trains very slowly (∼ 30 minutes for each cross-antibody model and ∼ 4 minutes for each per-antibody model on a single GPU), we only used it for the classification task in conjunction with the top-performing Antigen Res Mut embeddings.

**Figure S.1:**
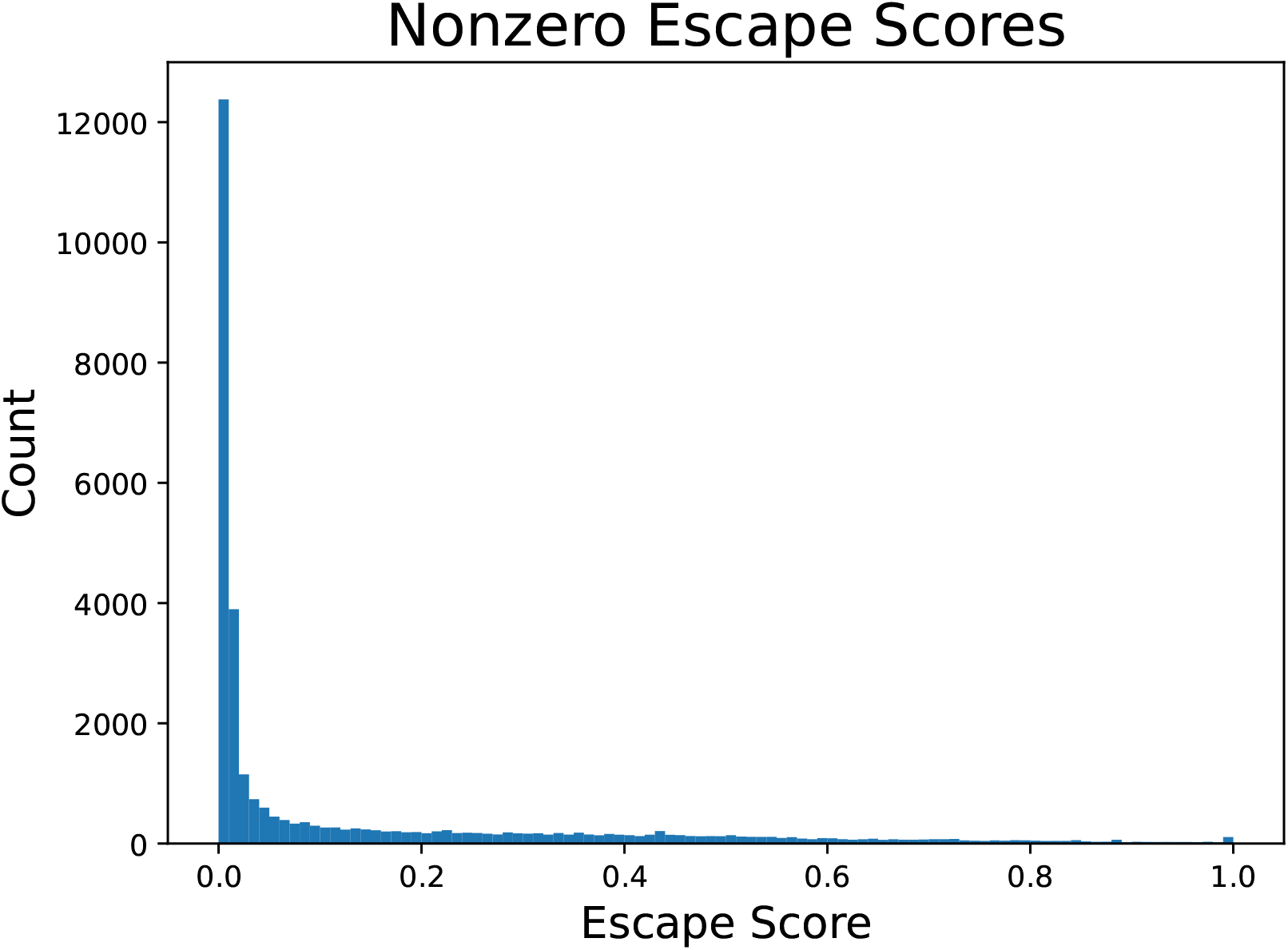
The distribution of the 30,658 non-zero escape scores in the SARS-CoV-2 deep mutational scanning data from Cao et al. [5]. Note: The 74 escape scores greater than 1 (max 3.6) are truncated to 1.

## D Complete Results

The remaining figures in the appendix show the complete set of results for all 168 combinations of data splits, models, and tasks that we ran. These results are also available in tabular form along with the data and embeddings at https://github.com/swansonk14/escape_embeddings.

**Figure S.2:**
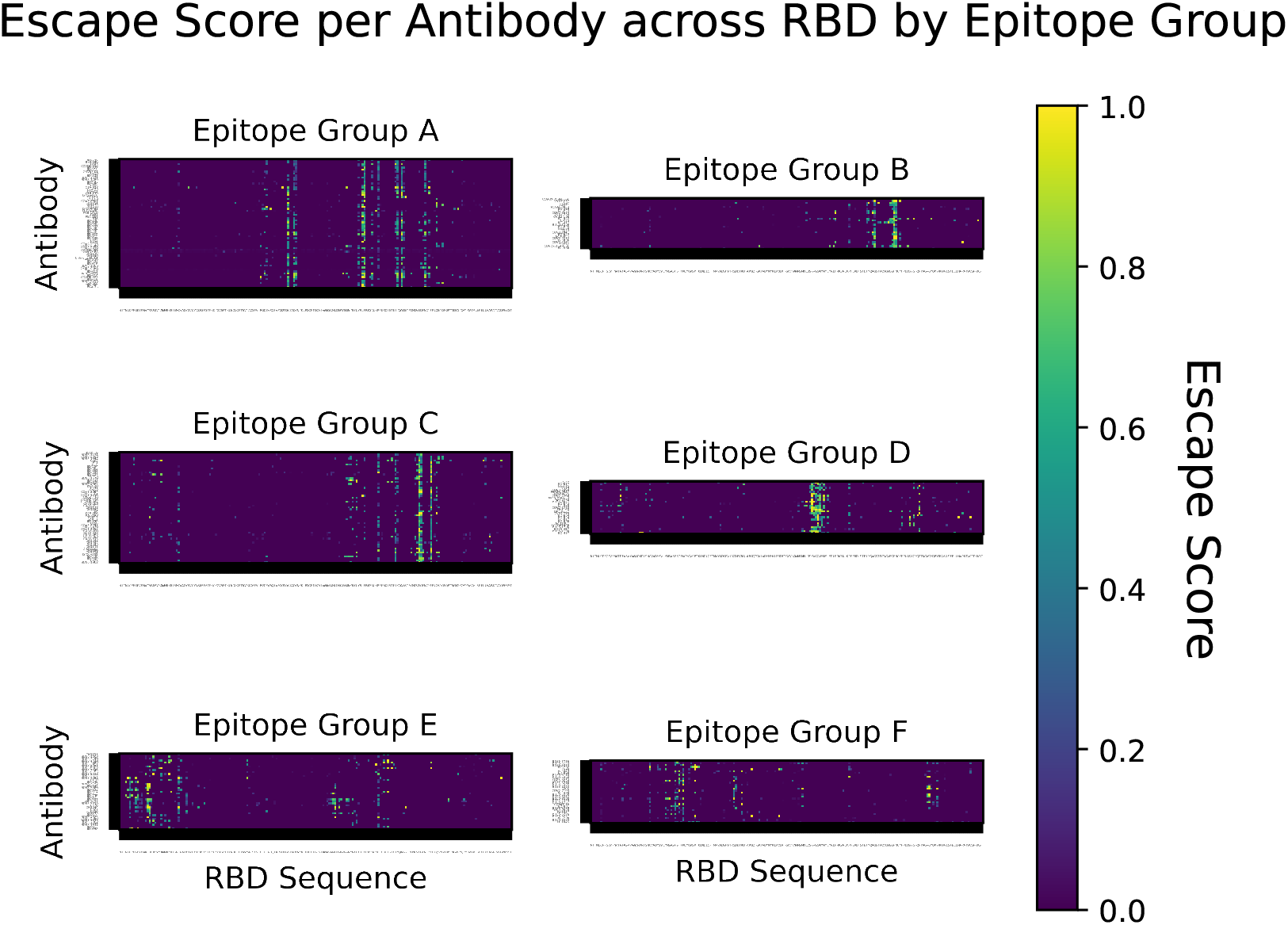
The average escape score across all amino acid mutations for each antibody and each antigen site in the receptor binding domain (RBD) of the SARS-CoV-2 spike protein, grouped according to the antibody clusters defined by Cao et al. [5]. Note: The 74 escape scores greater than 1 (max 3.6) are truncated to 1.

**Figure S.3:**
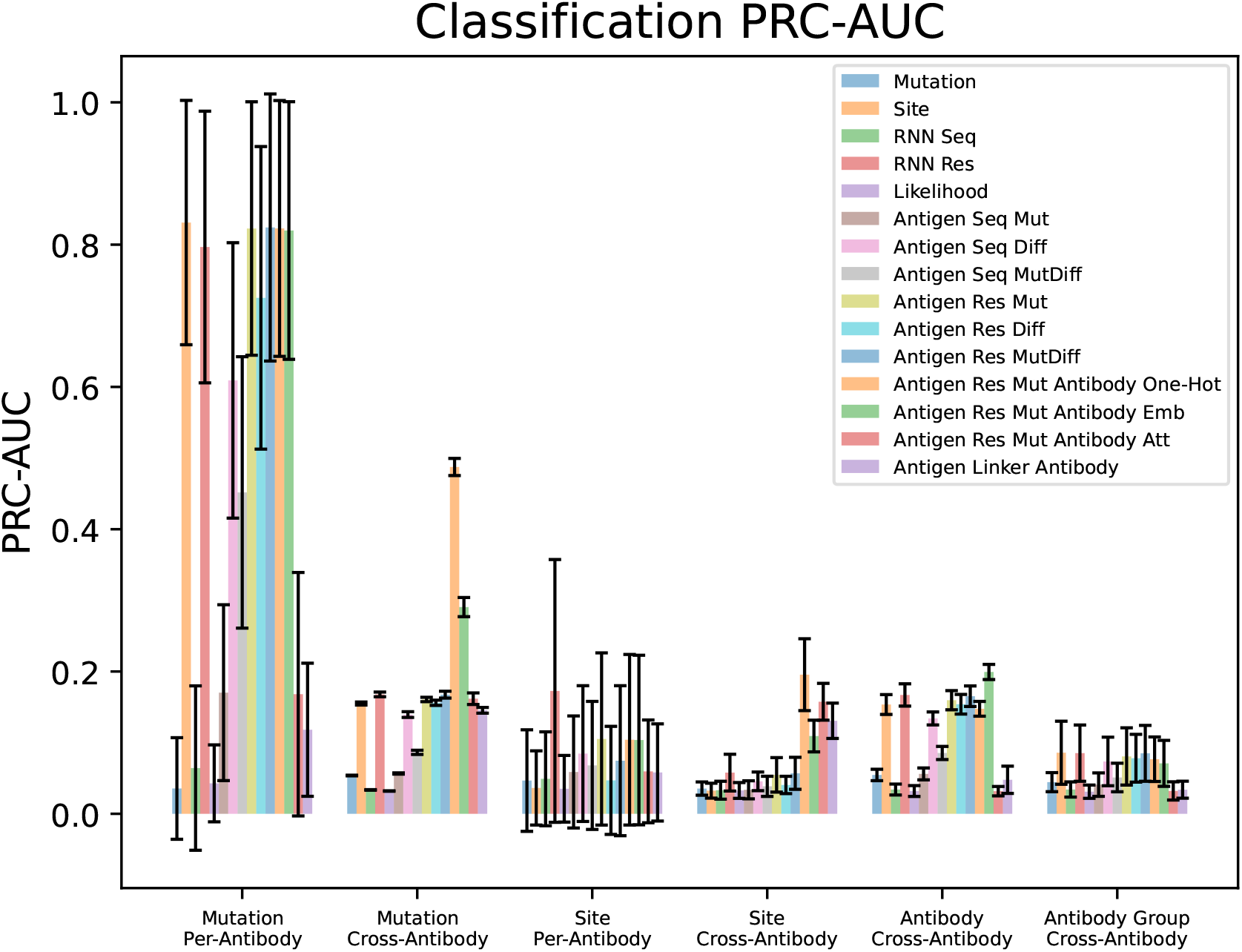
Classification model results using the PRC-AUC metric.

**Figure S.4:**
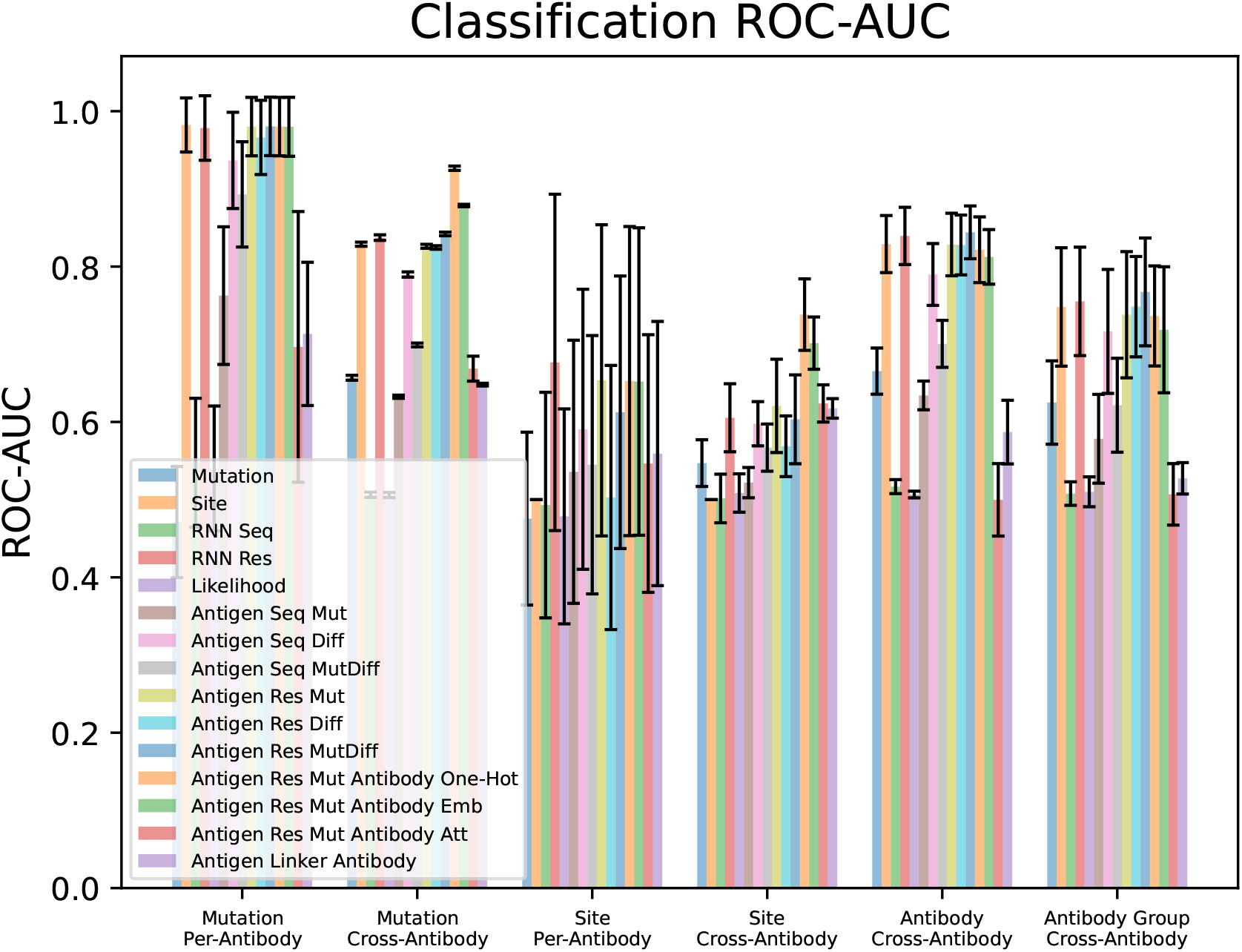
Classification model results using the ROC-AUC metric.

**Figure S.5:**
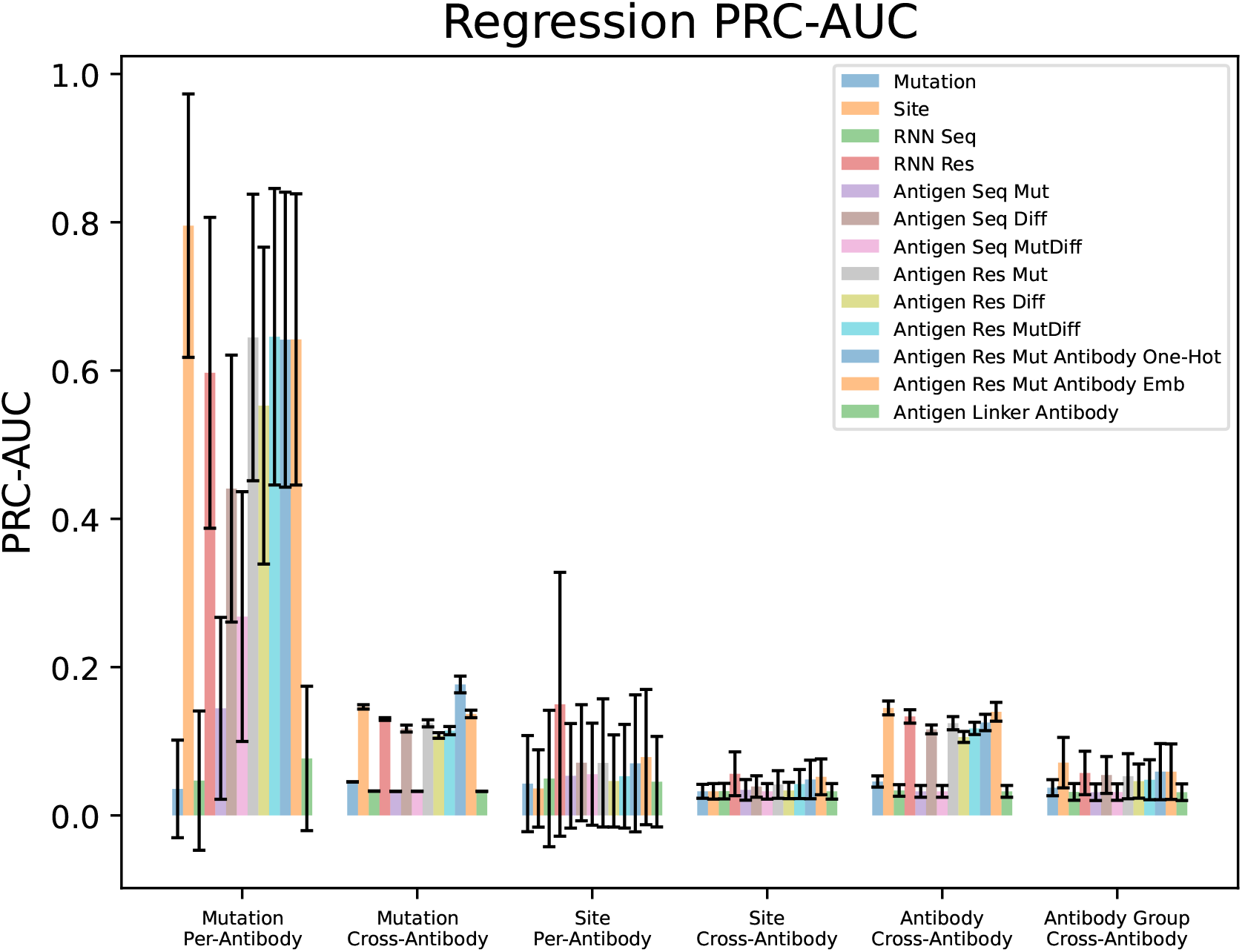
Regression model results using the PRC-AUC metric.

**Figure S.6:**
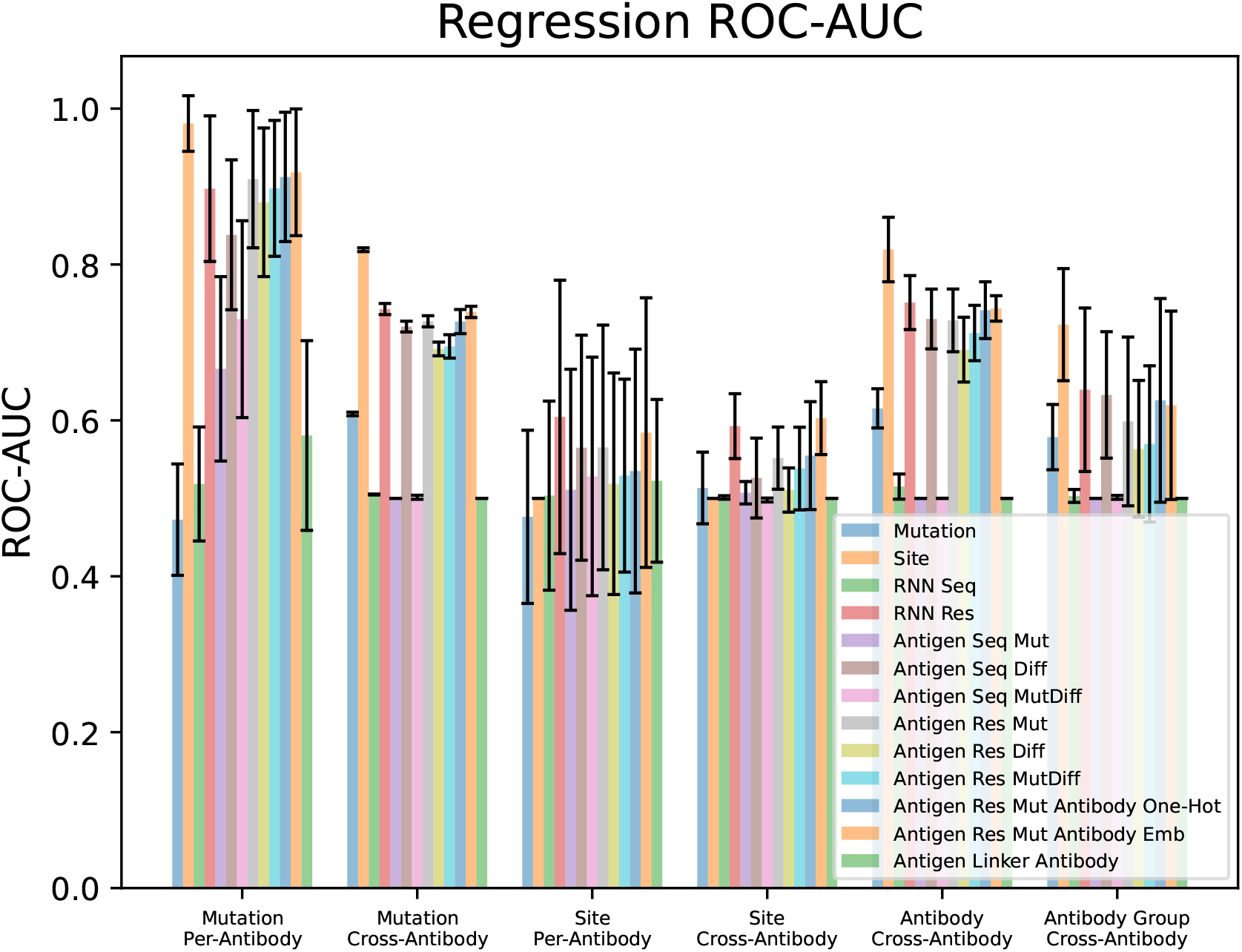
Regression model results using the ROC-AUC metric.

**Figure S.7:**
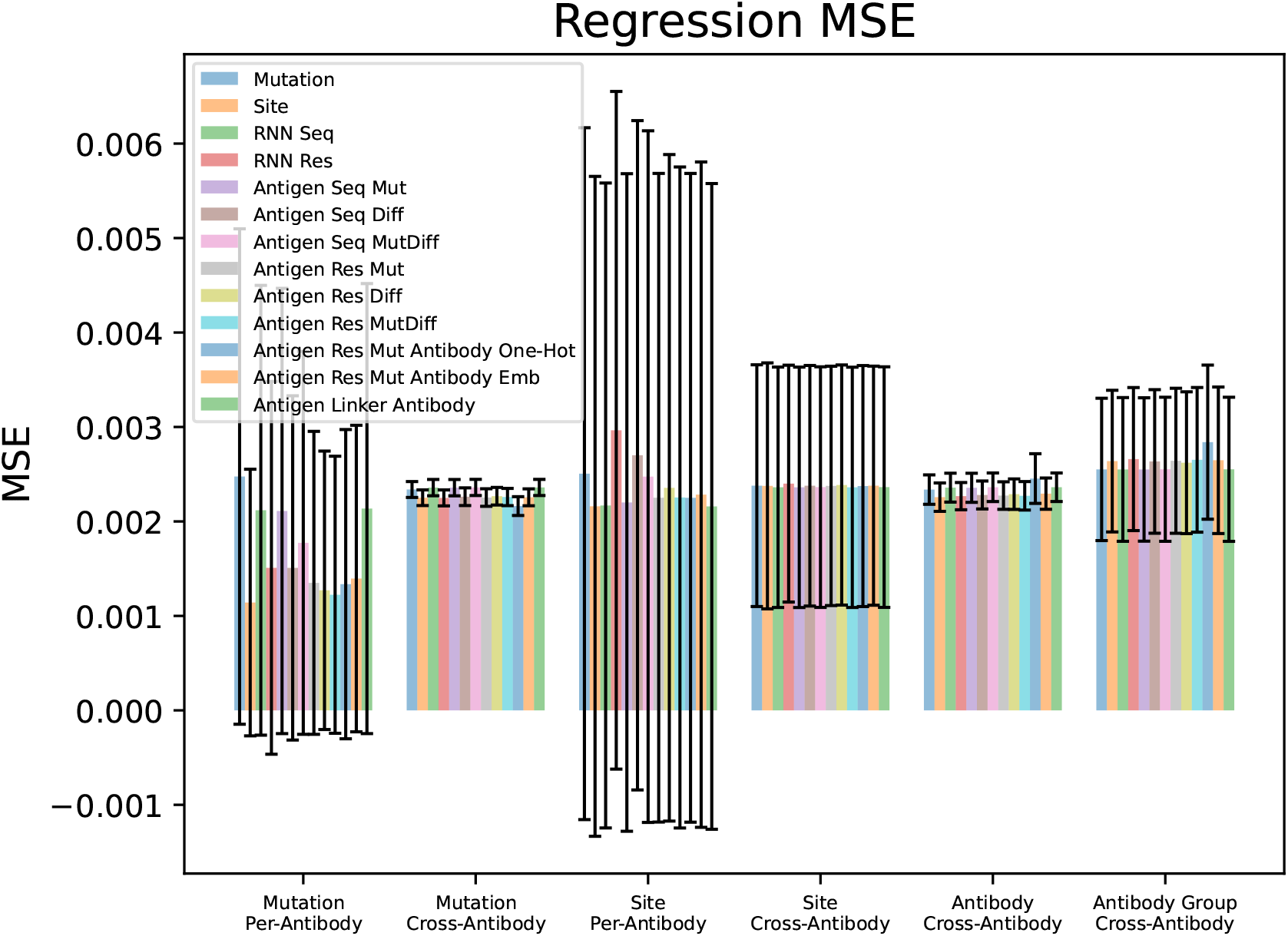
Regression model results using the MSE metric.

**Figure S.8:**
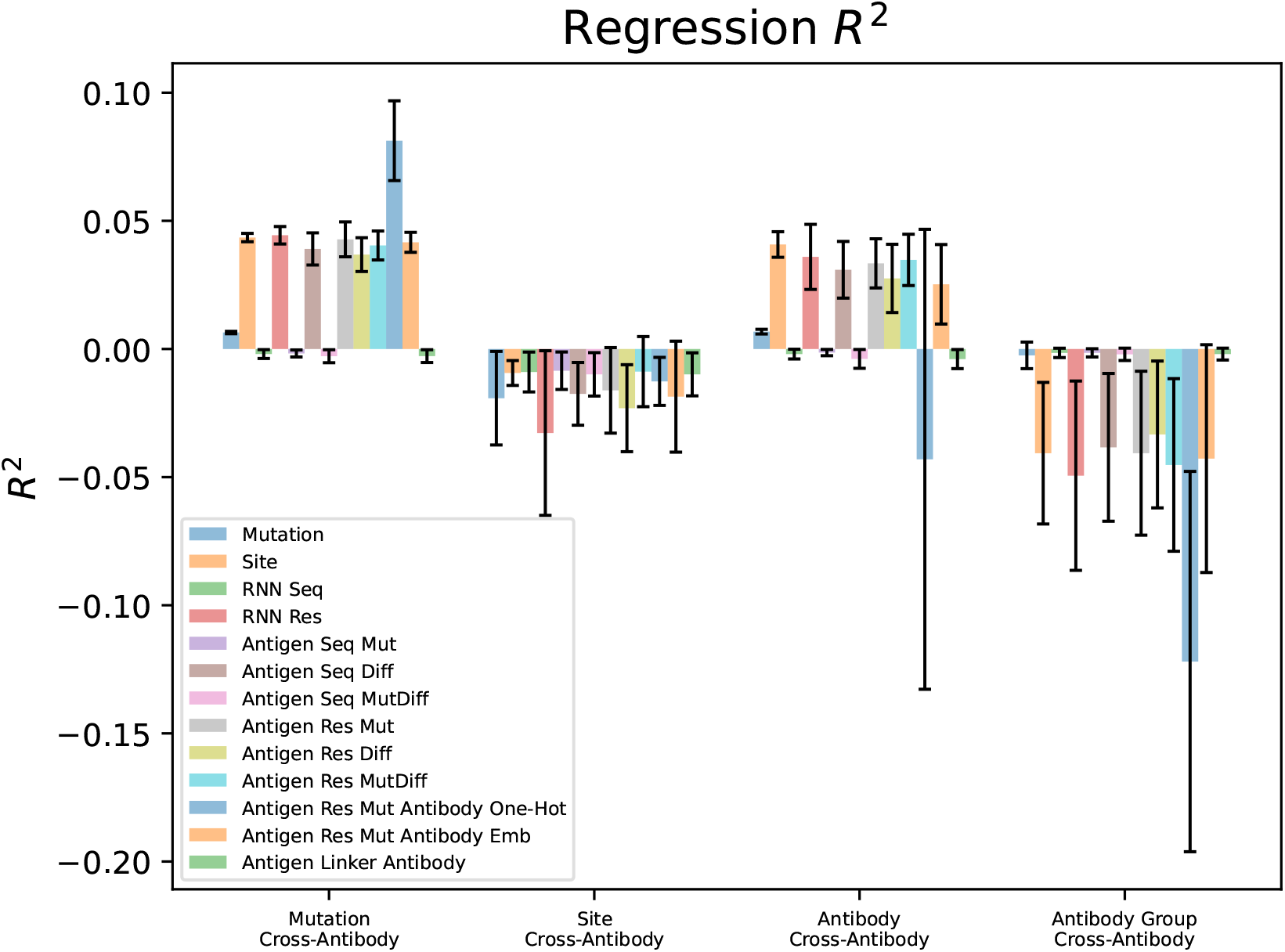
Regression model results using the R^2^ metric. Since the per-antibody splits have high variance, those splits are in separate plots below to make the scales legible in each plot.

**Figure S.9:**
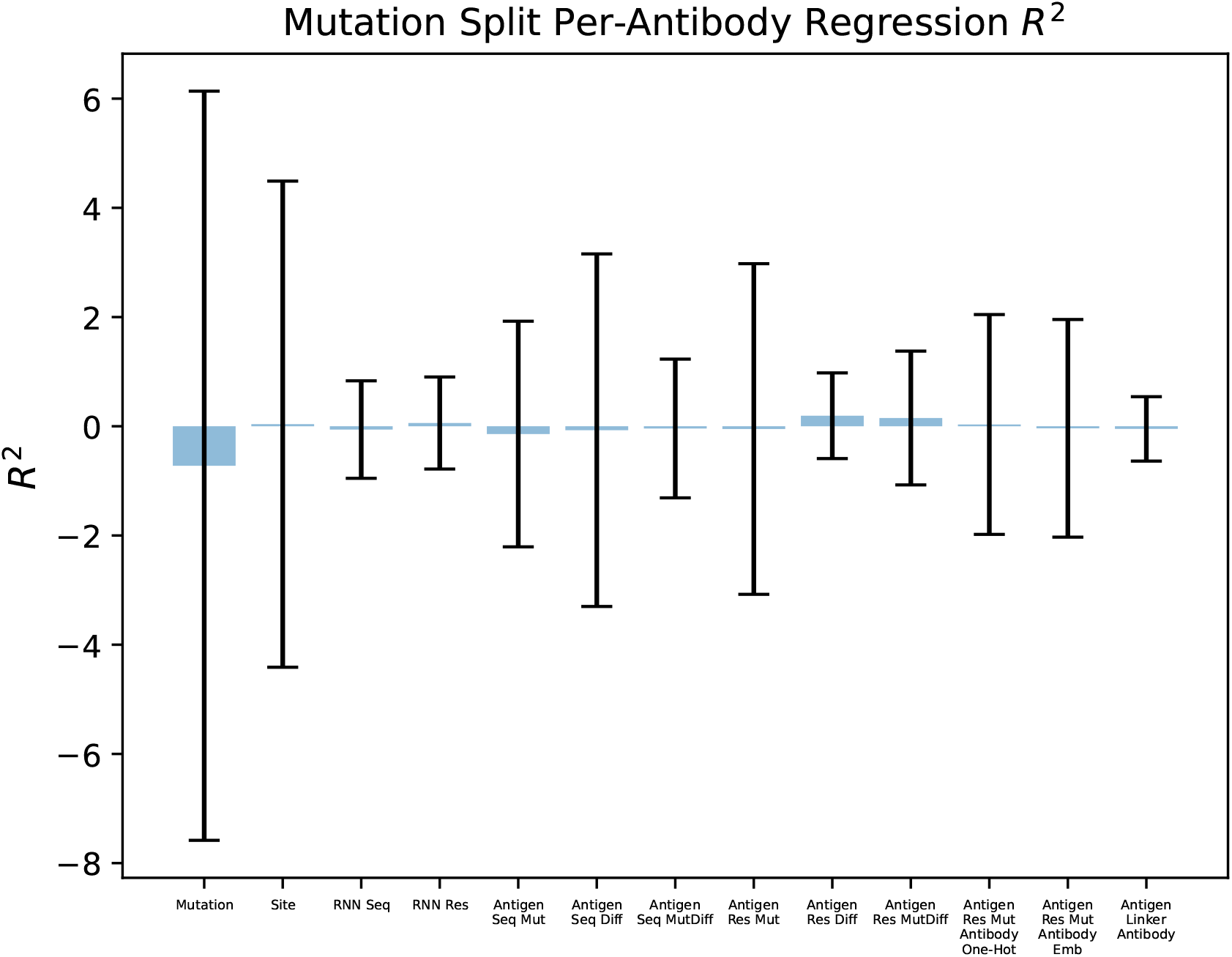
Regression model results using the R^2^ metric for the mutation per-antibody split.

**Figure S.10:**
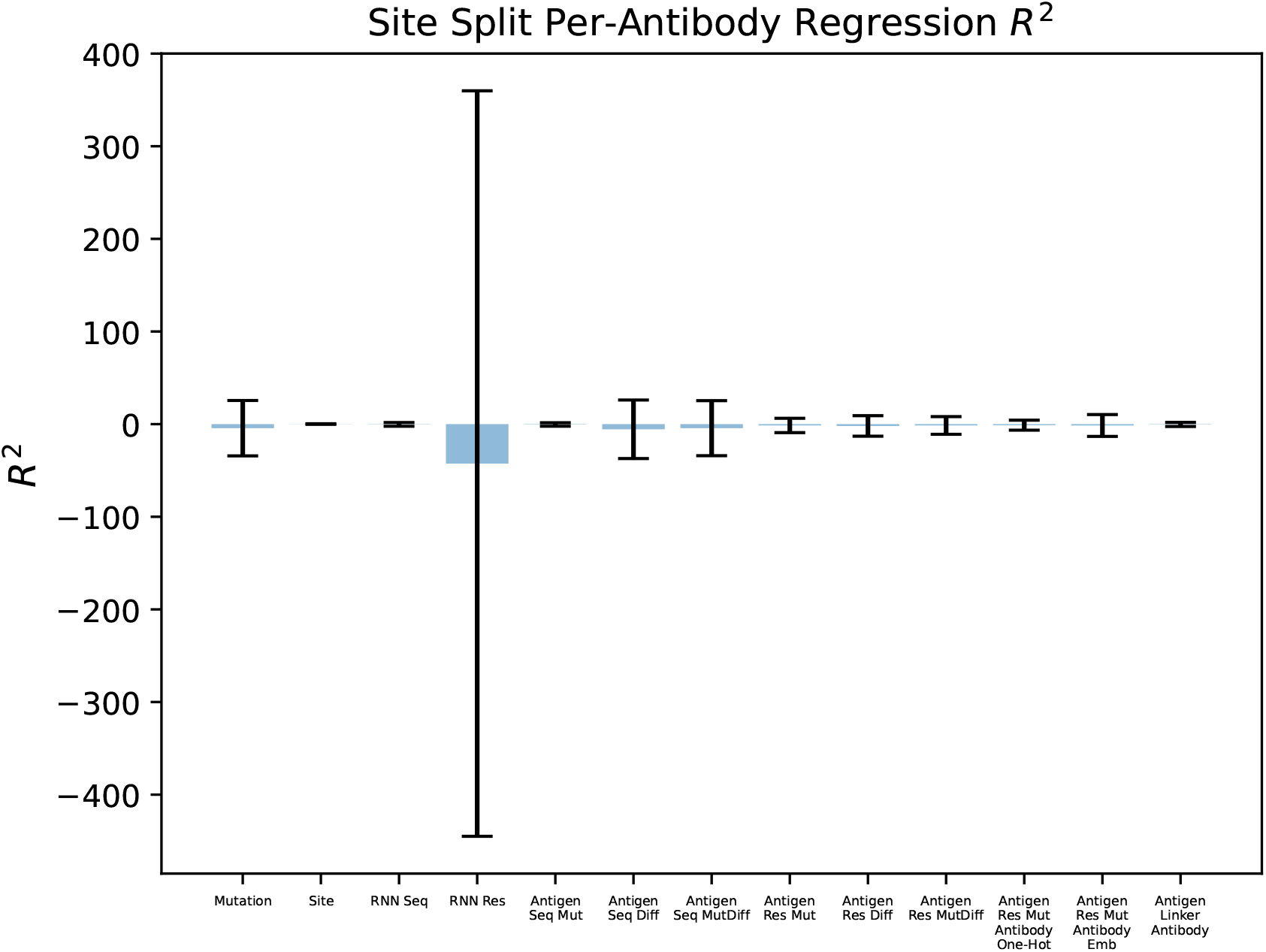
Regression model results using the R^2^ metric for the site per-antibody split.

## Footnotes

1 Our code, data, embeddings, and results are available at https://github.com/swansonk14/escape_embeddings

## References

1. Rai, K. R. et al. Acute Infection of Viral Pathogens and Their Innate Immune Escape. Frontiers in Microbiology 12. ISSN: 1664-302X. https://www.frontiersin.org/articles/10.3389/fmicb.2021.672026 (2021).

2. Kapingidza, A. B., Kowal, K. & Chruszcz, M. in Vertebrate and Invertebrate Respiratory Proteins, Lipoproteins and other Body Fluid Proteins (eds Hoeger, U. & Harris, J. R.) 465–497 (Springer International Publishing, Cham, 2020). ISBN: 978-3-030-41769-7. https://doi.org/10.1007/978-3-030-41769-7_19.

3. Starr, T. N. et al. Prospective mapping of viral mutations that escape antibodies used to treat COVID-19. Science 371, 850–854. eprint: https://www.science.org/doi/pdf/10.1126/science.abf9302. https://www.science.org/doi/abs/10.1126/science.abf9302 (2021).

4. Hie, B., Zhong, E. D., Berger, B. & Bryson, B. Learning the language of viral evolution and escape. Science 371, 284–288. eprint: https://www.science.org/doi/pdf/10.1126/science.abd7331. https://www.science.org/doi/abs/10.1126/science.abd7331 (2021).

5. Cao, Y. et al. Omicron escapes the majority of existing SARS-CoV-2 neutralizing antibodies. Nature 602, 657–663. ISSN: 1476-4687. https://doi.org/10.1038/s41586-021-04385-3 (Feb. 2022).

6. Frazer, J. et al. Disease variant prediction with deep generative models of evolutionary data. Nature 599, 91–95. ISSN: 1476-4687. https://doi.org/10.1038/s41586-021-04043-8 (Nov. 2021).

7. Meier, J. et al. Language models enable zero-shot prediction of the effects of mutations on protein function. bioRxiv. eprint: https://www.biorxiv.org/content/early/2021/11/17/2021.07.09.450648.full.pdf. https://www.biorxiv.org/content/early/2021/11/17/2021.07.09.450648 (2021).

8. Taft, J. M. et al. Deep mutational learning predicts ACE2 binding and antibody escape to combinatorial mutations in the SARS-CoV-2 receptor-binding domain. Cell. ISSN: 0092-8674. https://www.sciencedirect.com/science/article/pii/S0092867422011199 (2022).

9. Shan, S. et al. Deep learning guided optimization of human antibody against SARS-CoV-2 variants with broad neutralization. Proceedings of the National Academy of Sciences 119, e2122954119. eprint: https://www.pnas.org/doi/pdf/10.1073/pnas.2122954119. https://www.pnas.org/doi/abs/10.1073/pnas.2122954119 (2022).

10. Rives, A. et al. Biological structure and function emerge from scaling unsupervised learning to 250 million protein sequences. Proceedings of the National Academy of Sciences 118, e2016239118. eprint: https://www.pnas.org/doi/pdf/10.1073/pnas.2016239118. https://www.pnas.org/doi/abs/10.1073/pnas.2016239118 (2021).

11. Nijkamp, E., Ruffolo, J., Weinstein, E. N., Naik, N. & Madani, A. ProGen2: Exploring the Boundaries of Protein Language Models 2022. https://arxiv.org/abs/2206.13517.

12. Riesselman, A. J., Ingraham, J. B. & Marks, D. S. Deep generative models of genetic variation capture the effects of mutations. Nature Methods 15, 816–822. ISSN: 1548-7105. https://doi.org/10.1038/s41592-018-0138-4 (Oct. 2018).

13. Shin, J.-E. et al. Protein design and variant prediction using autoregressive generative models. Nature Communications 12, 2403. ISSN: 2041-1723. https://doi.org/10.1038/s41467-021-22732-w (Apr. 2021).

14. Lin, Z. et al. Language models of protein sequences at the scale of evolution enable accurate structure prediction. bioRxiv. eprint: https://www.biorxiv.org/content/early/2022/07/21/2022.07.20.500902.full.pdf. https://www.biorxiv.org/content/early/2022/07/21/2022.07.20.500902 (2022).

15. Hochreiter, S. & Schmidhuber, J. Long Short-Term Memory. Neural Computation 9, 1735– 1780. ISSN: 0899-7667. eprint: https://direct.mit.edu/neco/article-pdf/9/8/1735/813796/neco.1997.9.8.1735.pdf. https://doi.org/10.1162/neco.1997.9.8.1735 (Nov. 1997).

16. Evans, R. et al. Protein complex prediction with AlphaFold-Multimer. bioRxiv. eprint: https://www.biorxiv.org/content/early/2022/03/10/2021.10.04.463034.full.pdf. https://www.biorxiv.org/content/early/2022/03/10/2021.10.04.463034 (2022).

17. Paszke, A. et al. in Proceedings of the 33rd International Conference on Neural Information Processing Systems (Curran Associates Inc., Red Hook, NY, USA, 2019).

18. Suzek, B. E. et al. UniRef clusters: a comprehensive and scalable alternative for improving sequence similarity searches. Bioinformatics 31, 926–932. ISSN: 1367-4803. eprint: https://academic.oup.com/bioinformatics/article-pdf/31/6/926/569379/btu739.pdf. https://doi.org/10.1093/bioinformatics/btu739 (Nov. 2014).

19. Kingma, D. P. & Ba, J. Adam: A Method for Stochastic Optimization in 3rd International Conference on Learning Representations, ICLR 2015, San Diego, CA, USA, May 7-9, 2015, Conference Track Proceedings (eds Bengio, Y. & LeCun, Y.) (2015). http://arxiv.org/abs/1412.6980.

